# NECorr, a Tool to Rank Gene Importance in Biological Processes using Molecular Networks and Transcriptome Data

**DOI:** 10.1101/326868

**Authors:** Christophe Liseron-Monfils, Andrew Olson, Doreen Ware

## Abstract

The challenge of increasing crop yield while decreasing plants’ susceptibility to various stresses can be lessened by understanding plant regulatory processes in a tissue-specific manner. Molecular network analysis techniques were developed to aid in understanding gene inter-regulation. However, few tools for molecular network mining are designed to extract the most relevant genes to act upon. In order to find and to rank these putative regulator genes, we generated NECorr, a computational pipeline based on multiple-criteria decision-making algorithms. With the objective of ranking genes and their interactions in a selected condition or tissue, NECorr uses the molecular network topology as well as global gene expression analysis to find hub genes and their condition-specific regulators. NECorr was applied to Arabidopsis thaliana flower tissue and identifies known regulators in the developmental processes of this tissue as well as new putative regulators. NECorr will accelerate translational research by ranking candidate genes within a molecular network of interest.

## Introduction

In the last decade, functional genomics have been widely used in order to identify genes putatively involved in crop yield or response to stress. The differential expression of certain genes had served as first proof of their involvement in a particular biological process. Although these approaches could be successful (Hirai et al. 2007), they often represent only a partial snapshot of the molecular state within the plant. Consequently, understanding the regulation surrounding genes of interest for a biological process is difficult. Therefore, using only gene expression, it is extremely difficult to recreate the Gene Regulatory Network (GRN) underlying differential expression of the transcriptome. Finding computational approaches to this problem has led to the creation of the DREAM challenge (Schaffter, Marbach, and Floreano 2011). Most methods to study gene network rely on gene expression data as input. Then, gene networks are constructed and de-convoluted through co-expression analysis.

Among the different metrics to study co-expression networks, the most popular ones are the Pearson, Spearman, Kendall-Tau coefficients of correlation, or the mutual information. Some tools relying on these metrics were designed to study co-expression networks. Other tools relying on partial correlation have gained in popularity such as WGCNA (Langfelder and Horvath 2008). The difficulty with co-expression networks is the level of false positives due to non-direct gene interactions. The false positives are tentatively pruned away, but the pruning threshold remains a major problem.

Another approach to understanding the regulation of a biological process involves studying molecular networks. These networks serve two objectives: to identify gene modules linked to biological process regulation, and to predict the network behavior evolution under perturbations (Bansal et al. 2007). A protein-protein network (PPN) is based on the functional interaction between pairs of proteins. The yeast two hybrid system is one of the key techniques for identifying protein-protein interactions (Mohr and Koegl 2011). Databases recording experimentally validated or predicted PPN include BioGrid, Intact and the Arabidopsis interactome (Stark 2006; Hermjakob 2004; Wang et al. 2014).

A gene regulatory network (GRN) is a type of molecular network that relies on interactions between transcription factors and promoter sequences. Yeast one hybrid has been a method of choice to identify this type of interaction, notably in Arabidopsis (Reece-Hoyes and Walhout 2011). However, a new high throughput technique, enabling the genome-wide discovery of transcription factor binding sites was introduced recently (Reece-Hoyes and Walhout 2011; O’Malley et al. 2016); this technique is called DNA affinity purification sequencing (DAP-seq). DAP-seq doesn’t have the limitation of yeast-one hybrid, which is linked to the promoter tested in the yeast system. DAP-seq does not have the limitation of ChiP-seq that requires available antibodies for each tested transcription factor. DAP-seq is putatively able to produce a full GRN. The current version of the Arabidopsis cistrome resource encompasses ¼ of the transcription factors present in Arabidopsis (529/~2000).

Following the production of a network either using molecular or co-expression methodologies, the next step concerns the study of network dynamics. Methods for modeling such dynamics involve techniques such as machine learning algorithms, bayesian models or ordinary differential equations. Examples of ordinary differential equation methods are MNI using pre-existing expression ratios (di Bernardo et al. 2005), TSNI (Bansal, Della Gatta, and di Bernardo 2006) or NRI (Gardner et al. 2003). Once all possible perturbations are entered in the model; pre-existing gene expression ratios are used to predict future perturbations caused by one gene expression fluctuation in the network.

However, these models do not necessary help a researcher decide which genes may be the most important ones to act upon to modify a molecular process. To solve this problem, different prioritization algorithms were developed (Masoudi-Nejad et al. 2012; Lan et al. 2015). Most of the prioritization tools are linked to the medical field and the metrics to measure them are difficult to establish (Guala and Sonnhammer 2017). No clear methodology has emerged to apply ensemble methods aggregating the results of several methods (Kim, Farnoud, and Milenkovic 2015). Therefore, in order to analyze plant gene networks, there is a need for a methodology that takes into account available data and expert domain knowledge.

We generated a heuristic tool called NECorr to mine molecular network dynamics by taking advantage of transcriptomics data and network topology parameters. NECorr aims to prioritize the genes related to a biological phenomenon in order to enable researchers to perform further experimental validations. NECorr assigns weights to five different parameters reflecting their empirical importance.

Another aspect in network analysis is to identify edges of interest. In light of new advancements in genome editing, it becomes feasible to modify network edges such as transcription factor binding sites (Rodíguez-Leal et al. 2017). NECorr ranks the molecular network edges to prioritize the edges that would affect a biological process if disrupted by genome engineering or genetic crosses.

In the current analysis, NECorr was applied to define a ranked list of genes involved late flowering development. The Arabidopsis cistrome, derived from the DAP-seq analysis, was used as the molecular network source, in this case a GRN (Rodríguez-Leal et al. 2017; O’Malley et al. 2016). The Araport 11 Atlas was used as the transcriptome source (Rodríguez-Leal et al. 2017;O’Malley et al. 2016; Cheng et al. 2017). Known genes involved in flower development such as LEAFY (LFY) and GRXC7 (ROXY1) were found among the highest ranked genes in the results. However, new candidate genes without functional annotations were identified by the analysis.

## Materials and Methods

### Molecular network generation

The Gene Regulatory Network (GRN) coming from the Arabidopsis cistrome (Rodríguez-Leal et al. 2017; Cheng et al. 2017) was used as the molecular network to test NECorr. Using Araport11 gene annotation and the DAP-seq peaks, a GRN was constructed with edges corresponding to transcription factors binding from 1500 nt upstream to 500 nt downstream of gene transcription start sites (TSS). For genes with multiple transcripts, we selected the longest transcript with the longest translation to define the TSS. Within the molecular network, each node represents a product of a gene: transcript, mRNA or protein. This enables us to have a reduced and less complicated molecular network.

### Molecular network analysis

The molecular network was visualized using Cytoscape, yEd (Shannon et al. 2003). Different statistics were produced to capture the importance of each node within the molecular network. Centralities metrics such as eigenvector, betweenness, connectivity and pagerank centralities are part of those network statistics. Each centrality defines a different property that could be important for a scale free network. In the NECorr-Hub heuristic model, described below, the betweenness, the connectivity as well as the transitivity centralities are used. The betweenness centrality measures the number of shortest paths between in the network that pass through a given node. The connectivity is just the number of nodes linked to a particular node. The transitivity centrality defines the level of connection around a central node. It enables us to measure nodes generating an interconnected module around them.

### Transcriptome data

As starting material, two transcriptome datasets were used. The root spatio-temporal developmental atlas was applied to test and validate the NECorr model (Brady et al. 2007). The root data were collected by microarray analysis. Then, the developmental atlas of Arabidopsis from Araport 11 was used to find new regulators and important edges linked to specific tissues/conditions. This latter dataset is a compilation of 113 RNA-seq experiments (Cheng et al. 2017). We made the choice to use experiments not including mutant data or other Arabidopsis transgenic lines to avoid possible misinterpretations and biases caused by a possible alteration of the cell molecular network in these data.

### NECorr

The starting hypothesis of NECorr is that an important interaction for a stimuli response is a regulator acting on one or several hub genes. Hence, hub genes will propagate the systemic cascade appropriate to the stimuli. Thenceforth the dynamic of the molecular network will evolve. NECorr is composed of two steps to find the important interactions between a regulator and its hub gene.

### NECorr-Hub calculation

The first step is a heuristic model called NECorr-Hub which merges molecular network and gene expression data. It consists of ranking genes by importance within the molecular network as a function of gene co-expression across edges. The accepted inputs by NECorr are a molecular network and a gene expression table. Therefore, the ranking corresponds to the specific response of one condition compared to all other ones in the studied transcriptome data. NECorr-Hub is a linear model including 5 parameters: condition/tissue specificity of gene expression, co-expression of interactions across conditions, molecular network centralities betweenness, connectivity and transitivity. The importance given to each of these parameters was decided empirically.

Both genes of an interaction pair need to be co-expressed in most of the tissue/conditions showing that they can influence each other. Co-expression of genes in an interaction was considered as the most important parameter of the linear equation. In level of importance the second parameter was the gene expression specificity in the studied tissue/condition.

Finally, important genes for a network are hubs. These gene need to have a high level of connectivity in the molecular network to radiate on several genes to generate a proper response. The connectivity can be defined in several manners: betweenness, degree connectivity and transitivity were chosen as the most meaningful centralities to define gene importance in the hub.

After having decided of the level of importance of each parameter, each parameter weight was estimated using the Analytic Hierarchy Process (AHP) (Saaty 1977, 1987), a multiple-criteria decision analysis method. The AHP is applied through the R package pmr (“CRAN - Package Pmr” n.d.). The importance of the 5 parameters is generated by pairwise comparisons. Hence, this leads to an adjacency matrix of pairwise weight importance. From this adjacency matrix, Eigen-vectors are calculated to assign to a weight to each parameter. The AHP method is applied as follows. Each gene is ranked for the 5 parameters above. Each ranked parameter is standardized in values between 0 and 1 (z-score), in order to obtain data with the same scale. Then the weight is applied to each parameter value to obtain the final ranking value for each gene. For each tissue/condition, the parameter weights are applied as factors of a linear model that used to prioritize the gene importance.

For condition1:

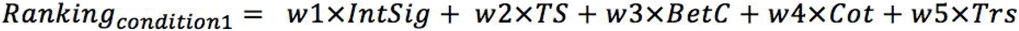

Where,

*IntSig: Interaction Significance (co-expression significance in the interaction involving the gene)*
*TS: Tissue specificity (selectivity)*
*BetC: Betweenness centrality*
*Cot: Connectivity centrality*
*Trs: Transitivity*

With the weights defined as follows:

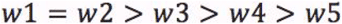

To rank each edge in the molecular network, the average ranking of the two genes defining this edge is taken to rank the interactions in the network for the condition. When several conditions are estimated, the gene ranking between conditions can be done by averaging each condition ranking.

### NECorr-Hub parameter estimation

Molecular network topology centralities are obtained using the R package iGraph (“CRAN - Package Igraph” n.d.). Co-Expression analysis (or the significance of each interaction) was estimated using a Rcpp script to evaluate the Gini correlation coefficient (GCC) related to each interaction. The GCC was previously shown to be an effective method for detecting transcription factor activity (Ma and Wang 2012). The co-expression significance for each gene is evaluated by averaging the magnitude of the correlation from all the interactions containing this particular gene using Fisher’s method (Mosteller and Fisher 1948; Poole et al. 2016).

The genes with tissue/condition specificity (or selectivity) are detected using the Intersection-Union test (IUT) with a relaxed threshold (raw p-value = 0.5) (“CRAN - Package Igraph” n.d.; Van Deun et al. 2009). The tissue/condition-selective genes or tissue/condition-excluded genes are assigned for each specific tissue/condition within a set of samples. This genes attributed to a tissue/condition are fuzzy due to the low selection threshold of the gene in IUT, therefore a gene could appear in a different tissue/condition as selective or excluded. Secondly, these selected genes are ranked for their tissue selectivity or exclusion using the Tissue Specificity index (Yanai et al. 2005. We define both a positive and negative TSI.

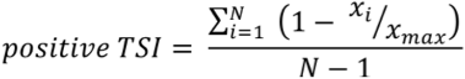

The negative Tissue specific index to measure the extent to which a gene is excluded from a tissue/condition.

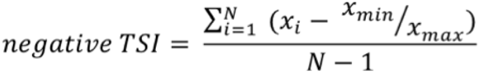

The results for TSI measurements are merged to obtain a ranking of all the tissue/condition selective/excluded genes defined from IUT test.

### NECorr effector

NECorr effector aims to discover regulators linked to the genes discovered by NECorr-Hub. Seven parameters were selected to try to find these regulators: the eigenvector centrality, the pagerank centrality, betweenness centrality, degree connectivity, gene differential expression, gene interaction significance and gene tissue selectivity. The pagerank and eigenvector centralities were added to find possible regulation as these centralities aim to discover nodes in a network linked to highly connected nodes. Several machine-learning algorithms were tested to discover effector genes. After testing on the Arabidopsis root data, the random forest algorithm was selected.

## Results and Discussions

### Explaining NECorr outputs

NECorr produces a series of outputs. The first type of output is the parameters used (Table S1). Each of the parameters used in the AHP algorithm is represented as a normalized value between 0-1.

The second output is the interaction correlation table with the metric used for the analysis, here the Gini correlation (Table S2).

The third output is the gene hub ranking (Table S3). The genes that can be important are ranked using the heuristic linear model from NECorr-Hub with the parameter weights calculated with the AHP. Then, each interaction present in the molecular network is ranked by averaging the hub rank of each of its two nodes or genes (Table S4). Finally, in the current version, NECorr offers an alternative to researchers to find genes influencing the molecular network. We called these genes effectors. NECorr effector aims to find targets of regulators. These targets may also be hub genes important a the studied biological processes that are found earlier with NECorr Hub (Table S5).

### Gene Ranking for flower

The top ranked gene was AT3G29080 (Table S3). Its individual parameter results show that the gene is highly tissue specific to the studied flower development (when compared the tissues present in the analysis - Table S1). The second gene has also an unknown function, AT1G15600. According to Gramene, this gene is part of an Arabidopsis specific tandem duplication of 7 genes relative to other Brassicas (Tello-Ruiz et al. 2018). Some of the other genes in this duplication are annotated as ubiquitin (AT1G15590, AT1G15625).

The results from the NECorr are shown to be not random as the 50 interactions with the highest rank are all interconnected showing that they are part of a putative biological process during the flower maturation at the bud stage 12 (Figure 1).

**Figure 1:**
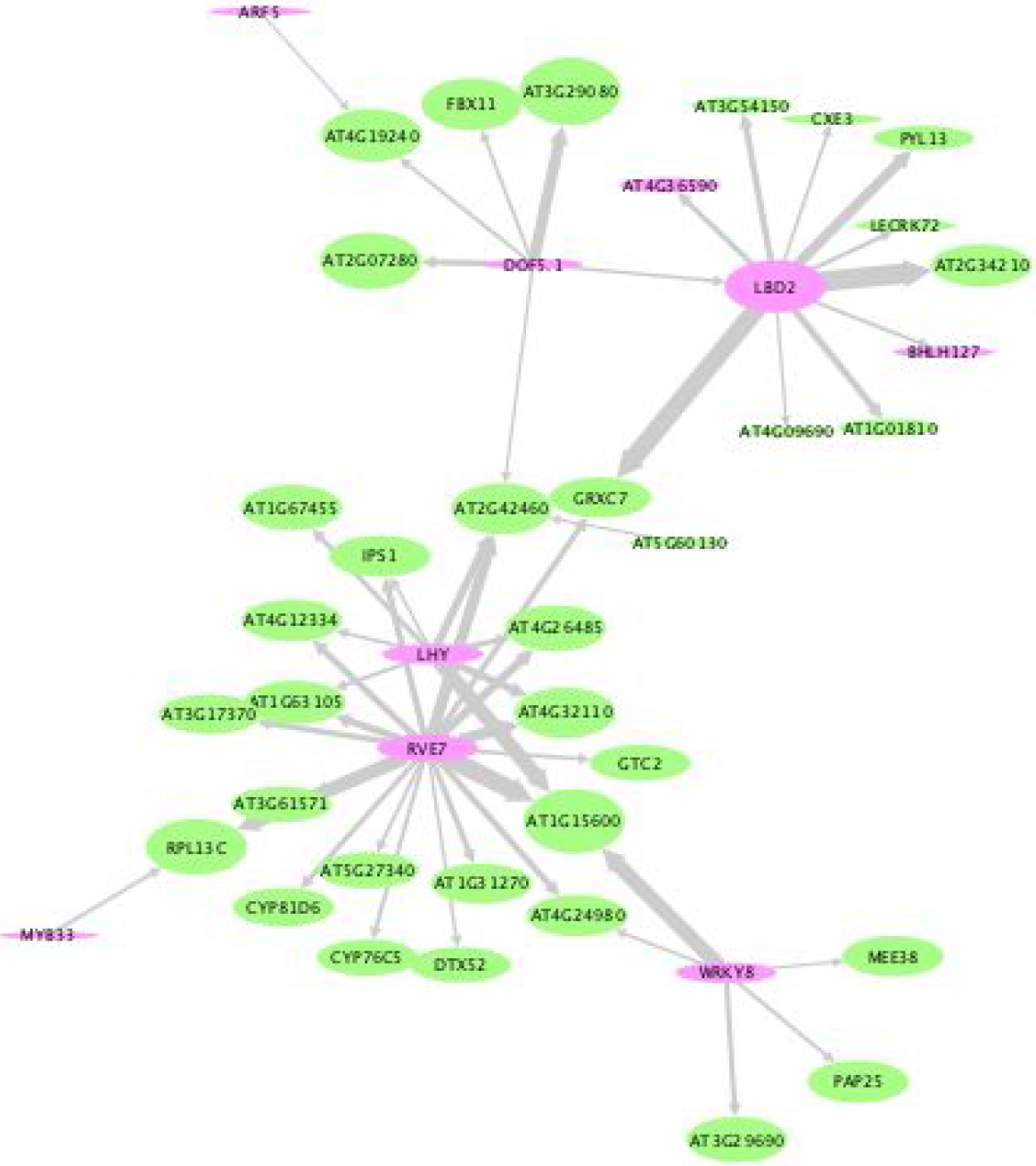
Sub-network of 50 highest ranked interactions by NECorr in the analysis of the floral bud stage 12. The size of the edges represents the significance of the ranked interactions. The size of the nodes is proportional to NECorr ranks. The green nodes represent the coding proteins; the pink nodes are the defined transcription factors.

The NECorr ranking of interactions (Table S3) found the best ranked interactions were LBD2 interacting with SPT5 and with GRXC7. GRXC7 (or ROXY1) is required during the development of petal (Xing, Rosso, and Zachgo 2005; Quon, Lampugnani, and Smyth 2017). Petal development occurs at the flower bud stage 12 explored in our analysis. GRXC7 is glutaredoxin influencing post-translational modifications of genes involved in petal development. Finally, these genes could be involved in the subtle balance occurring between layers during floral development, for instance anther development (Xing and Zachgo 2008). NECorr analysis of the cistrome implies that LBD2 could be one of the important regulators of GRXC7 during petal formation.

As expected, a NECorr analysis of the cistrome gives more insight into the regulation of genes already known to be involved in the late floral development. A more interesting result is to see the third ranked interaction in the analysis, RVE7 interacting with the promoter of AT1G15600. RVE7 is known for its role in the circadian rhythm (Li et al. 2011). However no function has been assigned to AT1G15600. The integrated analysis provided by NECorr could confer a role of this putative transmembrane protein in floral development.

From the NECorr analysis we could observe the interconnection of floral developmental genes and clock genes showing some possible regulation mechanisms between the light stimuli and the floral development.

### Limitations of NECorr

The concept of metagene where the gene node in the network could represent the mRNA, its protein, or its promoter give the possibility to bundle the gene form under one unique term: metagene (Moreau and Tranchevent 2012). The results are limited to the known metagene interactions that are within the molecular network. Consequently, the sparseness limits the possible discoveries. Furthermore, the putative function of the interactions can be discovered by the process of guilt-by-association to the studied biological process.

### Transfer to crop species

An Arabidopsis molecular network can be projected to a crop species via orthologous genes. Obviously, some of the interactions may be rewired if we compare molecular networks from 2 species; however an hypothesis is that the backbone of this network remains that same as a majority of the molecular processes are conserved. Theoretically, the orthologs of candidate genes found in Arabidopsis could be tested in a crop mutant lines for instance accelerating the discovery of putative lead genes linked to a biological process.

